# Th2 Cytokine Modulates Herpesvirus Reactivation in a Cell Type Specific Manner

**DOI:** 10.1101/2020.09.30.321778

**Authors:** Guoxun Wang, Christina Zarek, Tyron Chang, Lili Tao, Alexandria Lowe, Tiffany A. Reese

## Abstract

Gammaherpesviruses, such as Epstein-Barr virus (EBV), Kaposi’s sarcoma associated virus (KSHV), and murine γ-herpesvirus 68 (MHV68), establish latent infection in B cells, macrophages, and non-lymphoid cells, and can induce both lymphoid and non-lymphoid cancers. Research on these viruses has relied heavily on immortalized B cell and endothelial cell lines. Therefore, we know very little about the cell type specific regulation of virus infection. We have previously shown that treatment of MHV68-infected macrophages with the cytokine interleukin-4 (IL-4) or challenge of MHV68-infected mice with an IL-4-inducing parasite leads to virus reactivation. However, we do not know if all latent reservoirs of the virus, including B cells, reactivate the virus in response to IL-4. Here we used an *in vivo* approach to address the question of whether all latently infected cell types reactivate MHV68 in response to a particular stimulus. We found that IL-4 receptor expression on macrophages was required for IL-4 to induce virus reactivation, but that it was dispensable on B cells. We further demonstrated that the transcription factor, STAT6, which is downstream of the IL-4 receptor and binds a viral promoter in macrophages, did not bind to the viral promoter in B cells. These data suggest that stimuli that promote herpesvirus reactivation may only affect latent virus in particular cell types, but not in others.

**Importance:** Herpesviruses establish life-long quiescent infections in specific cells in the body, and only reactivate to produce infectious virus when precise signals induce them to do so. The signals that induce herpesvirus reactivation are often studied only in one particular cell type infected with the virus. However, herpesviruses establish latency in multiple cell types in their hosts. Using murine gammaherpesvirus-68 (MHV68) and conditional knockout mice, we examined the cell type specificity of a particular reactivation signal, interleukin-4 (IL-4). We found that IL-4 only induced herpesvirus reactivation from macrophages, but not from B cells. This work indicates that regulation of virus latency and reactivation is cell type specific. This has important implications for therapies aimed at either promoting or inhibiting reactivation for the control or elimination of chronic viral infections.

## Introduction

Herpesviruses establish chronic infections in their hosts. However, these viruses do not persistently replicate. Instead, they establish quiescent infections, termed latency. Latency is characterized by limited viral gene expression and no viral progeny production. Latent infections periodically reactivate and re-enter the lytic cycle to produce infectious virus. Latency and reactivation of herpesviruses is tightly controlled by the host immune system, reflecting the co-evolution of these viruses with their hosts. The development of drugs and vaccines to target herpesvirus, as well as efforts aimed at developing herpesviruses as vaccine vectors and oncolytic agents, requires that we understand the mechanisms that govern host control of herpesvirus latency.

The human γ-herpesviruses Epstein-Barr virus (EBV) and Kaposi’s sarcoma associated herpesvirus (KSHV) only infect humans and the cell culture systems available to study these viruses are limited to a few specific cell types. Pathogenesis and host-pathogen interaction of γ-herpesviruses can be examined using the close genetic relative of EBV and KSHV, murine gammaherpesvirus-68 (MHV68, γHV68). MHV68 establishes latency in macrophages and B cells and leads to lymphomas and tumors in immunocompromised mice, similar to EBV and KSHV in humans [1–6].

Because herpesvirus infections are ubiquitous, we previously examined the effects of co-infection on herpesvirus latency and reactivation. We showed that helminth parasites induce herpesvirus reactivation from latency. We identified the host cytokine, interleukin-4 (IL-4) and the transcription factor, signal transducer and activator of transcription (STAT)-6, that are required for parasite infection-driven reactivation of MHV68 in mice [7]. We determined that IL-4 receptor signaling through STAT6 and binding of STAT6 to a viral promoter induces MHV68 lytic gene expression. *In vitro*, treatment of bone marrow derived macrophages (BMDMs) with IL-4 increases virus replication. These data indicate that IL-4 can induce herpesvirus replication in macrophages *in vitro*, but it does not tell us which type of cells are responding to IL-4 *in vivo*.

Other host cytokines both positively and negatively regulate γ-herpesvirus latency in a cell-type-specific manner. IFNγ, for example, only suppresses MHV68 reactivation in the peritoneum from macrophages, but not in the spleen in B cells [8,9]. Similarly, IFNγ only suppresses KSHV lytic replication in certain cell types [10,11]. Other cytokines, including type I interferon and IL-21, also differentially impact MHV68 replication and latency in epithelial cells, macrophages, and B cells [12,13]. Understanding these cell-specific mechanisms will be critical for developing therapies for chronic viral infection that either enforce latency to prevent reactivation or promote reactivation to purge latent reservoirs and clear the virus.

In our previous work we detected robust IL-4-induced reactivation of MHV68 from peritoneal cells, but not from the spleen, perhaps indicating that IL-4 promotes MHV68 reactivation in a tissue and cell-type specific manner. To determine whether IL-4 promotes γ-herpesvirus reactivation in a cell type specific manner, we tested the requirements of IL-4 receptor (IL-4R) signaling in macrophages and B cells. We found that treatment of a latently infected B cell line with IL-4 did not lead to virus reactivation. Because of the limitations with *in vitro* systems, we examined the necessity of IL-4R signaling in macrophages and B cells *in vivo*. Using IL-4R-floxed mice crossed with either macrophage or B cell specific cre recombinase drivers, we determined the reactivation of MHV68 following treatment with IL-4/anti-IFNγ in mice lacking either IL-4R only on macrophages or only on B cells. We further examined the binding of STAT6 to a viral promoter of the latent-to-lytic switch gene (RTA, ORF50). Our results indicate that IL-4 induces herpesvirus reactivation and STAT6 binding to ORF50 promoter in a cell-type specific manner.

## Results

### IL-4 treatment increases viral replication in macrophages, but does not reactivate virus from latently infected B cell line

We first examined whether IL-4 reactivated MHV68 *in vitro* from two different cell types, macrophages and B cells, which harbor latent virus *in vivo*. In previous studies, we found that IL-4 treatment of bone marrow-derived macrophages (BMDMs) increased replication of MHV68 [7]. To confirm this, we treated BMDMs with IL-4 for 16 hours before infection with MHV68. As expected, IL-4 treatment increased the number of cells that express MHV68 lytic proteins, as measured by a flow cytometry assay, compared to untreated macrophages (Fig. 1A). To examine whether IL-4 has a similar effect on B cells as it does on macrophages, we took advantage of a B cell line (HE2.1), which harbors reactivation-competent latent MHV68 [14]. We treated the HE2.1 cells with IL-4 or Phorbol 12-myristate 13-acetate (PMA), a positive control that induces virus reactivation in B cells, and then examined expression of MHV68 lytic proteins by flow cytometry. While PMA induced viral reactivation from B cells, indicated by the increase in MHV68-positive cells, IL-4 treatment had no effect on MHV68 protein expression in the HE 2.1 B cells (Fig. 1B). We further measured virus-specific gene expression after IL-4 treatment in macrophages and B cells. In accordance with our flow cytometry data, IL-4 treatment increased expression of open reading frame (*Orf*) *50* and *Orf73* in infected macrophages (Fig. 1C). Although PMA treatment increased gene expression of *Orf50* and *Orf73* in HE2.1 B cells, we did not detect a significant increase of viral gene expression in this cell line after IL-4 treatment (Fig. 1D). These data suggest that IL-4-induced reactivation of MHV68 is cell type specific. However, there are caveats with the use of cell lines and *de novo* infection of primary B cells with MHV68 in tissue culture is limited. We therefore examined IL-4-induced reactivation from specific cell types *in vivo*.

**Figure 1.**
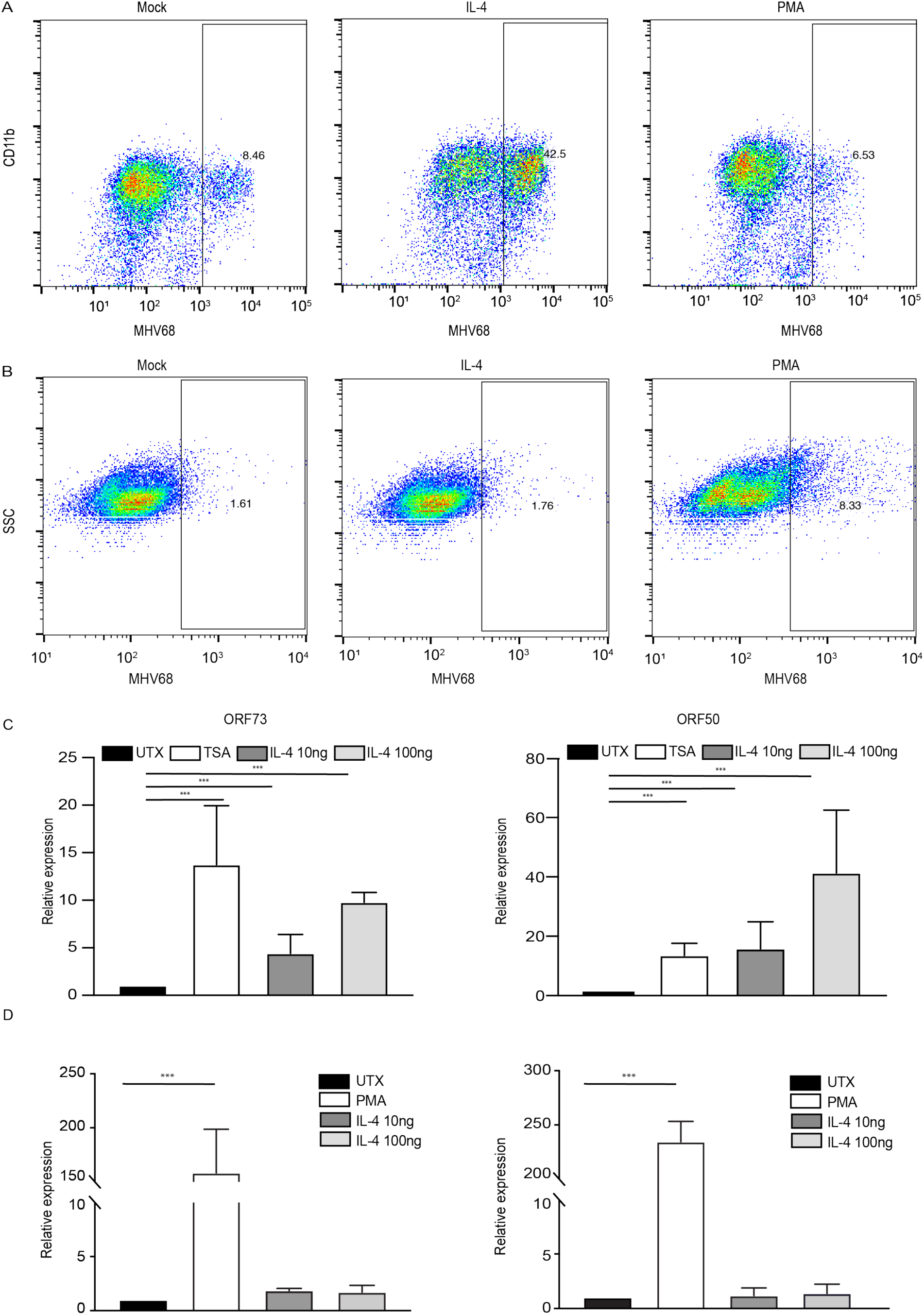
IL-4 treatment increases viral replication in macrophages but does not reactivate virus from latently infected B cell line. A. Bone marrow-derived macrophages (BMDMs) were treated with 100 ng/ml IL-4 and 20 ng/ml PMA for sixteen hours in culture medium and then infected with MHV68 at MOI=5. Twenty-four hours after infection, cells were fixed and cells expressing lytic viral proteins were determined by flow cytometry. B. HE2.1 B cells were treated with 100 ng/ml IL-4 and 20 ng/ml PMA for 38 hours, then cells were fixed and cells expressing lytic viral proteins were determined by flow cytometry. C. BMDMs were treated with 130 nM TSA or different concentrations of IL-4 for sixteen hours and then infected with MHV68 at MOI=5. Transcripts of virus genes *Orf50* and *Orf73* were determined at 12 hours after infection. Expression was normalized to *Gapdh*. Data are from three independent experiments. D. HE2.1 B cells were treated with 20 ng/ml PMA or different concentrations of IL-4 for 24 hours. qRT-PCR was conducted to assess expression of virus genes *Orf50* and *Orf73*. Expression was normalized to *Gapdh*. Data are from three independent experiments.

### STAT6 is not required for MHV68 lytic replication

Before examining the role of IL-4 signaling in macrophages and B cells during latency, we first tested whether IL-4 signaling was required for control of acute MHV68 replication. We compared virus replication in wildtype mice or *Stat6*^*-/-*^ mice, because *Stat6*^*-/-*^ mice are deficient in signaling downstream of the IL-4R*α*. After infecting mice with MHV68, spleen samples were collected on days two, four, and seven post infection. Viral titers were determined by plaque assay. The *Stat6*^-/-^ mice had similar levels of viral replication compared to wildtype mice, indicating that STAT6 is not essential for MHV68 acute replication (Fig 2).

**Figure 2.**
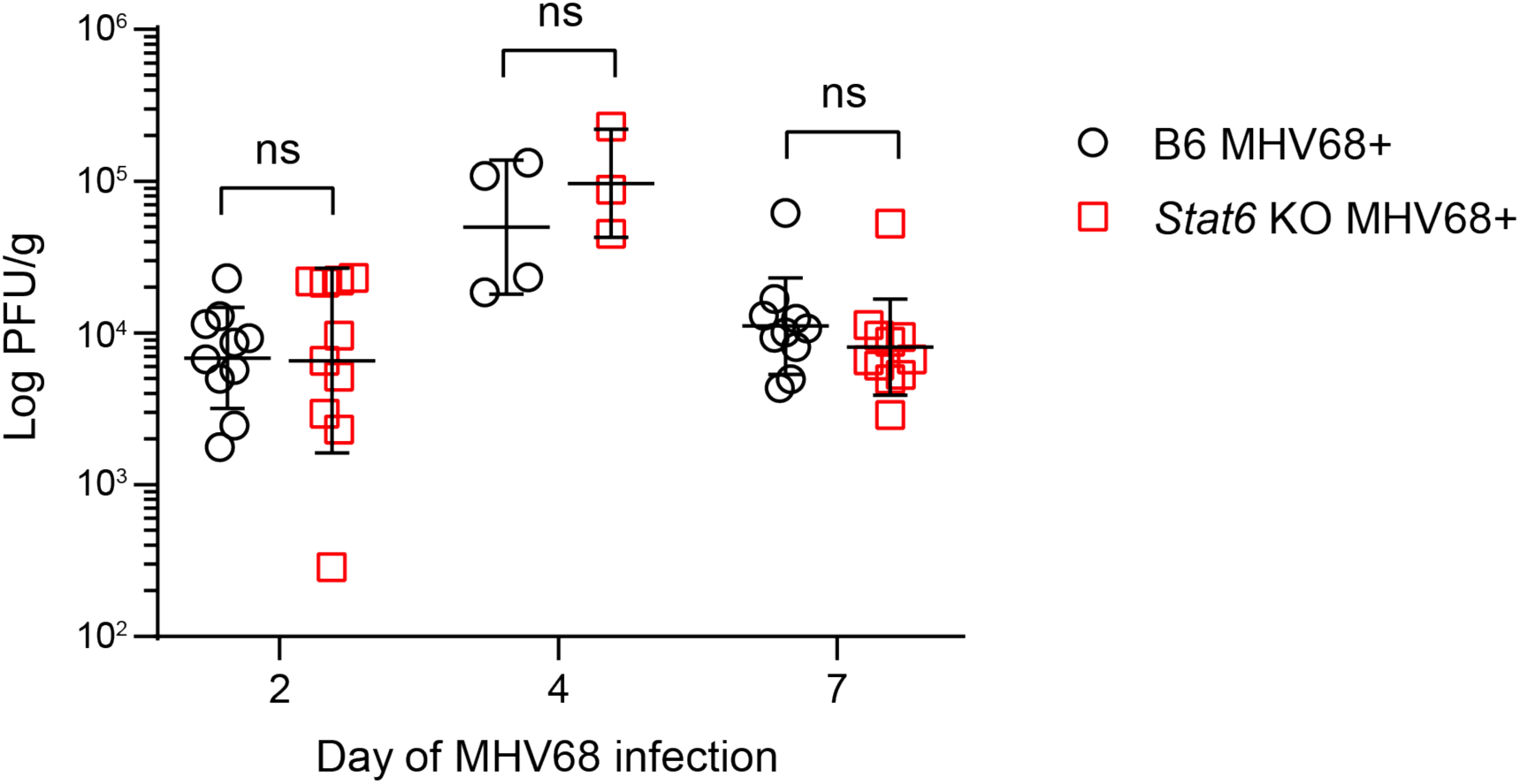
STAT6 is not required for MHV68 replication in the spleen. Mice were infected by intraperitoneal injection with 10^6^ PFU of MHV68. Whole spleens were collected and snap frozen at day 2 (D2), day 4 (D4) and day 7 (D7) post-infection to quantitate lytic replication of MHV68. Virus titer was determined by plaque assay. Bars represent mean ± SD. Each dot represents an individual mouse. ns, not significant.

### IL-4 receptor expression is required for MHV68 reactivation from macrophages

To determine whether IL-4-induced viral reactivation *in vivo* requires IL-4 receptor (IL-4R) on macrophages, we generated mice that are deficient of IL-4R specifically on myeloid cells, including macrophages. We crossed mice homozygous for loxP-flanked *Il4rα* genes, *Il4rα flox/flox* (*Il4rα*^*f/f*^) [16] with mice that express cre recombinase under the control of the lysozyme-M (*LysM*) promoter (*LysMcre*). Gene knockout efficiency in *Il4rα*^*f/f*^ *x LysMcre+ mice* was demonstrated by the absence of detectable levels of IL-4R expression in macrophages from peritoneal exudate cells (PECs) with and without virus infection (Fig 3A and S1A). B cell expression of IL-4R in the peritoneum was normal in the *Il4rα*^*f/f*^ *x LysMcre+* mice (Fig. 3A).

**Figure 3.**
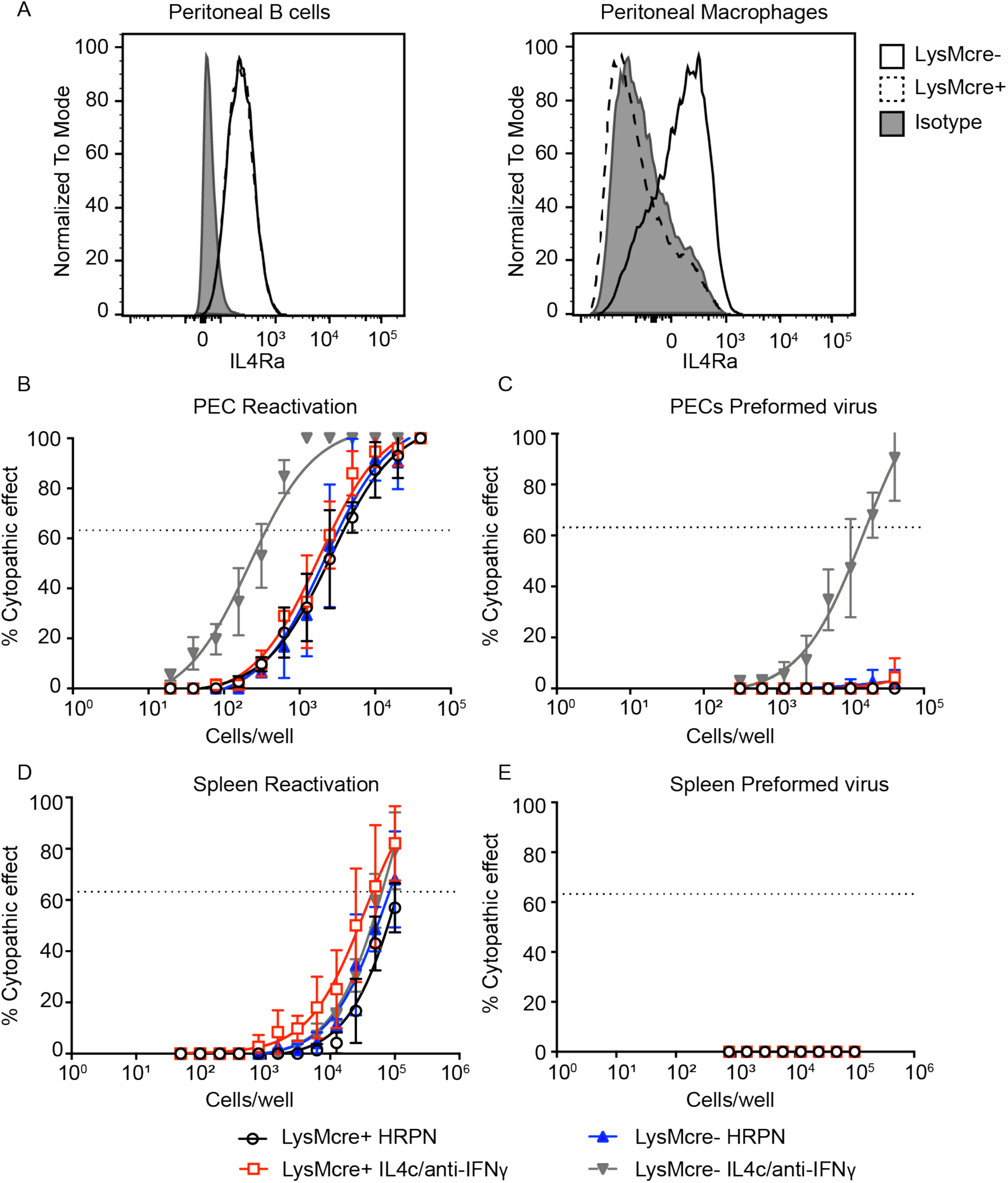
IL-4R expression is required for MHV68 reactivation from macrophages. A. Peritoneal exudate cells (PECs) were collected from uninfected *Il4rα*^*f/f*^ *x LysMcre+* and *Il4rα*^*f/f*^ *x LysMcre-* mice and flow cytometry was used to detect the IL-4 receptor on peritoneal B cells and macrophages. Representative histograms of two independent experiments are shown. B-E. *Il4rα*^*f/f*^ *x LysMcre+* and *Il4rα*^*f/f*^ *x LysMcre-* mice were intraperitoneally infected with 10^6^ PFU wild-type MHV68. Mice received isotype control (HRPN) or both IL-4c and anti-IFNγ after 28 days post-infection. Another dose of IL-4c was administered two days after the first treatment and then all mice were euthanized at 5 days posttreatment. Peritoneal exudate cells (PECs) (B and C) and splenocytes (D and E) were processed into single-cell suspensions. Reactivation frequencies (B and D) were determined by *ex vivo* plating of serially diluted PECs and splenocytes on a MEF monolayer. Presence of preformed virus in PECs and splenocytes was determined by disrupting the cell suspensions and plating serially diluted samples on MEF monolayers (C and E). Cytopathic effect was scored 3 weeks post-plating. Groups of 3-5 mice were pooled for each experiment. Data represents the average of 3 independent experiments. The error bars represent standard errors of the mean.

We previously showed that injection with long-lasting IL-4 complexes (IL-4c) combined with blocking antibody to interferon (IFN)-γ (IL-4c/anti-IFNγ) mimicked intestinal helminth infection to induce MHV68 reactivation. While virus reactivation induced by parasite infection is highly variable, reactivation induced by IL-4c/anti-IFNγ is more robust, demonstrated by both bioluminescence imaging and an *ex vivo* reactivation assay [7]. Therefore, we chose to use IL-4c/anti-IFNγ to examine cell-type specific reactivation *in vivo*.

To determine if IL-4R expression is required on macrophages for virus reactivation *in vivo*, we infected *Il4rα*^*f/f*^ *x LysMcre* positive and negative mice with MHV68. To induce reactivation from latency, we injected mice with either isotype control (HRPN) or IL-4c/anti-IFNγ one month after infection. Seven days after treatment, peritoneal cells, which harbor latent virus primarily in macrophages and B cells [4,17], were collected for a limiting dilution assay (LDA) to quantitate virus reactivation. In this *ex vivo* reactivation assay, explanted cells are plated on a mouse embryonic fibroblast (MEF) monolayer in 96-well plates in a limiting dilution fashion. Cytopathic effect (CPE) is detected after three weeks, and the frequency of reactivating cells is determined using Poisson’s distribution [18]. We can distinguish *ex vivo* reactivating virus from *in vivo* preformed virus by plating both live explanted cells and lysed cells. The lysed samples (termed disrupted) induce cytopathic effect of the MEF monolayers if they contain virus that reactivated *in vivo* prior to collection of samples. The live explanted cells induce cytopathic effect of the MEF monolayers when virus reactivates during *ex vivo* culture. Similar to our previous findings, when *Il4rα*^*f/f*^ *x LysMcre*-mice are injected with IL-4c/anti-IFNγ, we observed substantial virus reactivation from latency compared with isotype-injected mice (HRPN) (Fig. 3B). Moreover, we detect preformed virus in the *Il4rα*^*f/f*^ *x LysMcre-* mice injected with IL-4c/anti-IFNγ (Fig. 3C), indicating that virus reactivated *in vivo* prior to collection. In contrast, when *Il4rα*^*f/f*^ *x LysMcre*+ mice, which lack IL-4R on macrophages, were injected with IL-4c/anti-IFNγ, there was no increase in virus reactivation *ex vivo* and no preformed virus (Fig. 3B, C). These data indicate that macrophage expression of IL-4R is required for IL-4c/anti-IFNγ induced virus reactivation in PECs. We also measured virus reactivation from splenocytes, which harbor latent virus primarily in B cells [19] and detected no significant increase in virus reactivation or preformed virus from either *LysMcre+* or *LysMcre-* mice injected with IL-4c/anti-IFNγ (Fig. 3D, E).

Defective reactivation could be due to a failure of MHV68 to establish or maintain latency in IL-4R deficient macrophages. To test this, the frequency of virus-positive peritoneal cells during chronic infection was quantitated by limiting dilution nested PCR targeting MHV68 *Orf72* [18]. Equivalent numbers of viral genomes were detected in *Il4rα*^*f/f*^ *x LysMcre+* and *Il4rα*^*f/f*^ *x LysMcre-* mice (Fig. S1B), indicating that latency was established at equivalent levels in *Il4rα*^*f/f*^ *x LysMcre+* and *Il4rα*^*f/f*^ *x LysMcre*-mice.

To further confirm our LDA reactivation results, *Il4rα*^*f/f*^ *x LysMcre+* and *Il4rα*^*f/f*^ *x LysMcre-* mice were infected with luciferase tagged MHV68, MHV68-M3FL [20] and injected with IL-4c/anti-IFNγ. Mice were imaged five days after injection in accordance with our previously published results [7]. As expected, *LysMcre*-mice that received IL-4c/anti-IFNγ displayed increased reactivation, whereas *LysMcre*+ mice did not, verifying that IL-4-induced reactivation *in vivo* requires IL-4R expression on macrophages (Fig. S2). Collectively, these data suggest that macrophage expression of IL-4 receptor is required for IL-4c/anti-IFNγ induced virus reactivation *in vivo*.

### IL-4R expression on macrophages is not required for Cyclosporin A induced reactivation

Our results raise the possibility that loss of IL-4 receptor expression on macrophages may impact the ability of MHV68 to reactivate not just to IL-4 stimulation, but also to other reactivation agents *in vivo*. To test if *in vivo* reactivation is functional in *Il4rα*^*f/f*^ *x LysMcre* mice, we administered cyclosporin A (CA) intraperitoneally. Cyclosporin A is an immunosuppressive drug that reactivates MHV68 [20,21]. It targets multiple pathways in T cells that inhibit activation and proliferation. It is believed that cyclosporin A increases reactivation of MHV68 because T cells, especially CD8+ T cells, are important for control of MHV68 [22,23]. Reactivation induced by cyclosporin A should occur through IL-4 receptor independent pathways.

*Il4rα*^*f/f*^ *x LysMcre*+ and *Il4rα*^*f/f*^ *x LysMcre*-mice were infected with MHV68-M3-FL, to track virus reactivation *in vivo*. Infected mice were injected with cyclosporin A on day 40 and day 42 of infection (Fig. 4A). The mice were imaged at several timepoints to capture the peak of reactivation, which occurred day 7 post cyclosporin A treatment (Fig. 4B, C). We detected reactivation from both *LysMcre*+ and *LysMcre*-mice after cyclosporin A treatment (Fig. 4C), indicating that lack of the IL-4 receptor does not inhibit *in vivo* reactivation of MHV68 in response to cyclosporin A.

**Figure 4.**
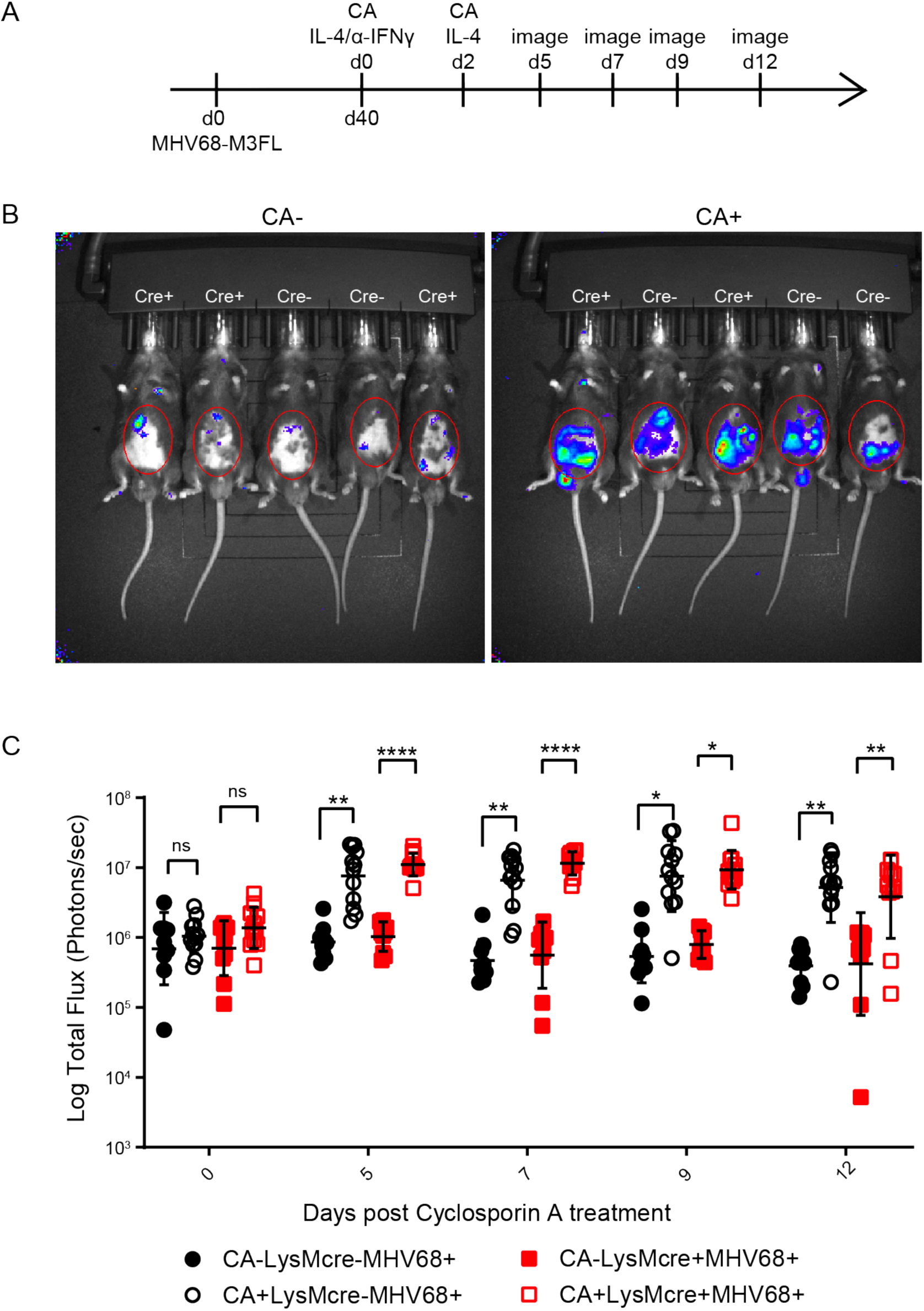
IL-4R expression on macrophages is not required for Cyclosporin A induced reactivation. A. Timeline of treatments with Cyclosporine A (CA) and imaging to measure CA-induced MHV68-M3-FL reactivation in mice. Total flux (photons/second) was measured using IVIS bioluminescence imager. B. Representative images of *Il4rα*^*f/f*^ *x LysMcre+* and *Il4rα*^*f/f*^ *x LysMcre-* mice on day 7 post CA treatment. The red circles mark the area used for total flux measurements. C. Quantitation of total flux. Data shown were the results obtained from a pool of two independent experiments. Bars represent mean ± SD. Each dot represents an individual mouse. ns, not significant; * P<0.05, ** P<0.01, **** P<0.001.

### The absence of IL-4 receptor on B cells does not affect IL-4c/anti-IFNγ induced MHV68 reactivation

Since MHV68 also establishes latency in B cells, we next evaluated MHV68 reactivation frequency in mice lacking IL4R*α* on B cells. We crossed *Il4ra*^*f/f*^ mice with *Cd21-cre* expressing mice to delete B cell expression of IL-4R*α*. We used *Cd21-cre* rather than the more commonly used *Cd19-cre* because both IL-4R*α* and Cd19 are on the same mouse chromosome. Cd21 is expressed on germinal center B cells and B1b cells in the peritoneum, both of which are latency reservoirs of MHV68 [24–26]. As shown in Figure 5A, we used flow cytometry to confirm that cre recombinase successfully deleted the IL-4 receptor on B cells in the spleen and peritoneum of *Il4rα*^*f/f*^ *x Cd21-cre+* mice (Fig. 5A). IL-4R expression is intact in *Il4rα*^*f/f*^ *x Cd21-cre*+ macrophages (Fig. 5A).

**Figure 5.**
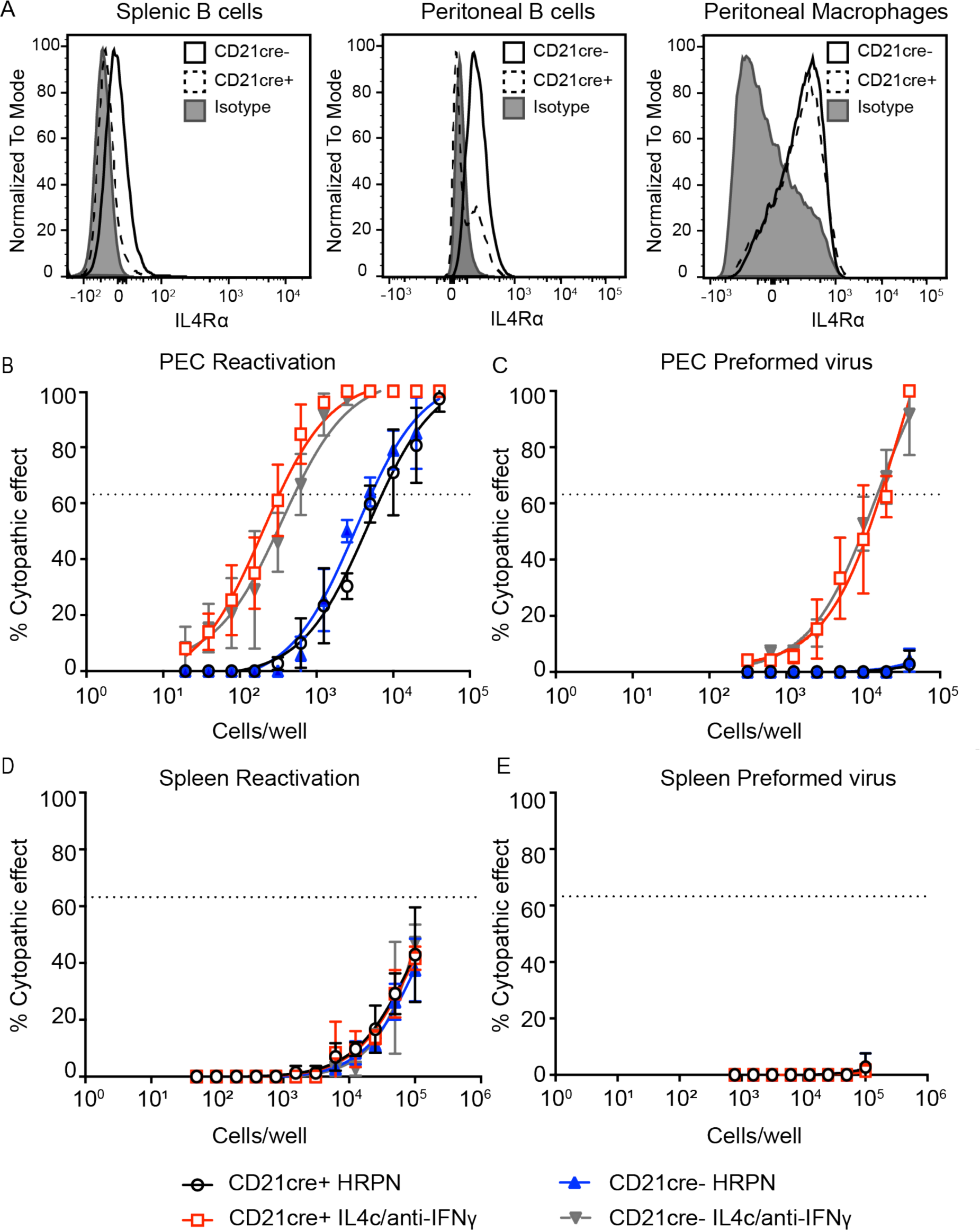
IL-4R expression is not required for MHV68 reactivation from B cells. A. Whole spleen or peritoneal exudate cells (PECs) were collected from uninfected *Il4rα*^*f/f*^ *x Cd21-cre+* and *Il4rα*^*f/f*^ *x Cd21-cre-* mice, and flow cytometry was used to detect IL-4R on B cells and macrophages. Representative histograms of two independent experiments are shown. B-E. *Il4rα*^*f/f*^ *x Cd21-cre+* and *Il4rα*^*f/f*^ *x Cd21-cre-* mice were intraperitoneally infected with 10^6^ PFU wild-type MHV68. Mice received isotype control (HRPN) or both IL-4c and anti-IFNγ 28 days post-infection. Another dose of IL-4c was administered two days after the first treatment and then mice were euthanized 5 days post-treatment. Splenocytes (D and E) and PECs (B and C) were processed into single-cell suspensions. Reactivation frequencies (B and D) were determined by *ex vivo* plating of serially diluted PECs and splenocytes on a MEF monolayer. Presence of preformed virus in PECs and splenocytes was determined by disrupting the cell suspensions and plating serially diluted samples on MEF monolayers (C and E). Cytopathic effect was scored 3 weeks post-plating. Groups of 3-5 mice were pooled for each infection and analysis. The data are pooled from 3 independent experiments. The error bars represent standard errors of the mean.

We assayed the frequency of viral reactivation from *Il4rα*^*f/f*^ *x Cd21-cre+* and *Il4rα*^*f/f*^ *x Cd21-cre-* mice with and without IL-4c/anti-IFNγ in the peritoneum and in the spleen. As expected, injection of latently infected *Il4rα*^*f/f*^ *x Cd21-cre-* mice with IL-4c/anti-IFNγ induced virus reactivation from explanted peritoneal cells (Fig. 5B) and induced virus reactivation *in vivo*, as detected by the presence of preformed virus (Fig. 5C). Similar to *Cd21-cre-* mice, *Il4rα*^*f/f*^ *x Cd21-cre+* mice injected with IL-4c/anti-IFNγ also reactivated the virus to comparable levels as cre-recombinase negative mice (Fig. 5B, C). In summary, our results demonstrate that the absence of IL-4R on B cells in the peritoneum did not dampen MHV68 reactivation upon IL-4c/anti-IFNγ treatment.

We also examined virus reactivation from the spleen following IL-4c/anti-IFNγ treatment. We did not detect an increase in virus reactivation from the spleen after IL-4c/anti-IFNγ treatment from either group of mice (Fig. 5 D, E), indicating that IL-4c/anti-IFNγ does not stimulate splenic reactivation of MHV68.

### STAT6 does not bind the promoter of RTA/ORF50 in B cells

We showed previously in macrophages that phosphorylated STAT6 directly binds to a viral promoter, termed N4/N5, of *Orf50*/RTA which is the latent-to-lytic switch gene [7,27]. Activation of *Orf50* promotes viral replication *in vitro* in macrophages and virus reactivation *in vivo* [28–32]. Despite the fact that B cells in the germinal centers respond to IL-4 produced by T follicular cells [33], one possibility is that IL-4 stimulation of B cells does not activate STAT6. We determined if IL-4 signaling was intact in the HE2.1 B cells by measuring phosphorylation of STAT6. We treated the B cells with low dose and high dose of IL-4 and probed for phosphorylation of STAT6. As shown in the Fig 6A, both doses of IL-4 treatment induced phosphorylation of STAT6, indicating that the HE2.1 B cells indeed respond to IL-4.

**Figure 6.**
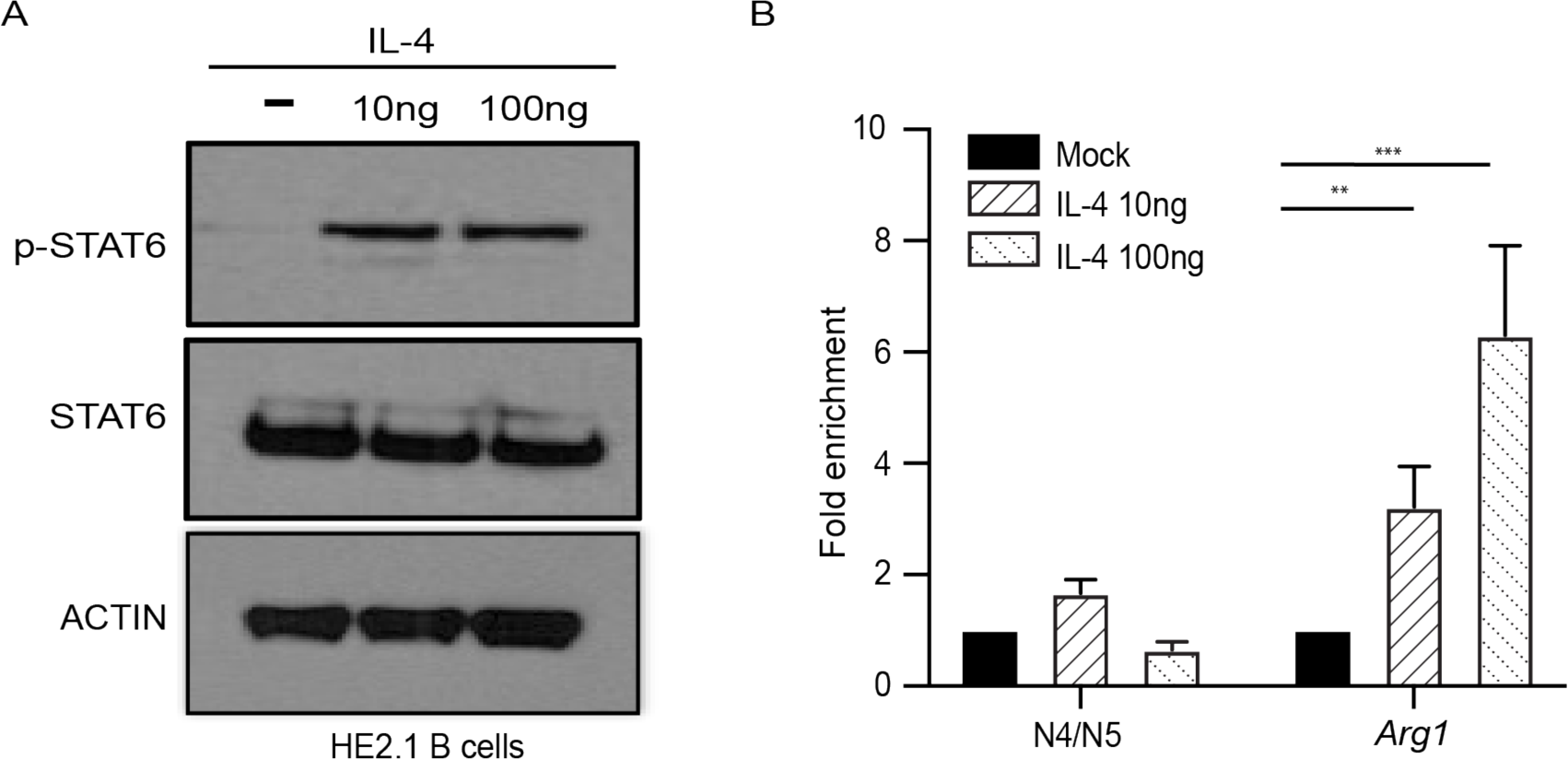
STAT6 does not bind the promoter of RTA/*Orf50* in HE2.1 B cells. A. HE2.1 B cells were treated with different concentrations of IL-4 for 24 hours. Total and phosphorylated STAT6 proteins were determined by western blotting. B. HE2.1 B cells were treated with different concentrations of IL-4 for 24 hours. DNA bound to STAT6 was immunoprecipitated, and quantitative PCR (qPCR) was performed for the N4/N5 promoter region or Arginase-1 *(Arg1*). Percentage of input after normalizing to immunoglobulin G control was calculated for both *N4/N5* and *Arg*1. Data are from three independent experiments. Error bars are standard error of the mean. P-values calculated by one-way ANOVA. ** P<0.01, **** P<0.001

Another possibility is that STAT6 does not bind to the N4/N5 promoter of *Orf50/*RTA in B cells. To test this, we determined STAT6 binding to the *Orf50*/RTA promoter N4/N5 in B cells. Using STAT6 antibody-based ChIP-qPCR, we examined STAT6 binding affinity in HE2.1 B cells after IL-4 treatment. Compared to increased binding to the *Arg1* promoter, which is activated by STAT6 signaling, STAT6 did not bind to the N4/N5 promoter of viral *Orf50/*RTA upon IL-4 stimulation of HE2.1 B cells (Fig. 6B). These data suggest that even though IL-4 activates STAT6 in B cells, STAT6 does not bind to the viral *Orf50* promoter in B cells.

## Discussion

We found that expression of IL-4 receptor on macrophages was required for efficient MHV68 reactivation from latency upon administration of IL-4c plus blocking antibody to IFNγ. In contrast, reactivation was normal in response to Cyclosporin A in mice with IL-4R deficiency on macrophages. Intriguingly, IL-4 receptor deficiency on B cells did not impair virus reactivation in response to IL-4c combined with anti-IFNγ treatment. We also revealed that even though IL-4 treatment of both macrophages and B cells induces phosphorylation of STAT6 in both cell types, phosphorylated STAT6 does not bind to the ORF50 promoter in B cells. Thus, our study suggests that different latently infected cell populations respond differentially to reactivation signals.

For gammaherpesviruses, reactivation from latency is controlled by one or more master regulators of transcription. Expression of these latent-to-lytic switch genes is silenced during latency, and reactivation signals turn on expression of these genes. When expressed, these transactivator proteins induce expression of lytic viral genes, leading to production of virus. In the case of MHV68, this master regulator is *Orf50* (also called RTA) [34]. We demonstrated previously that IL-4 treatment of macrophages leads to induction of *Orf50* expression. We found that the host transcription factor downstream of the IL-4R, STAT6, binds to one of the promoters of *Orf50* in macrophages [7]. In this report we found that even though IL-4 induces phosphorylation of STAT6 in B cells, STAT6 did not bind to *Orf50* N4/N5 promoter. These data suggest that this promoter is inaccessible to STAT6 binding in B cells and may indicate that the *Orf50* N4/N5 promoter is differentially accessible to transcription factor binding in B cells and macrophages.

Interestingly, there are multiple *Orf50* promoters for MHV68, termed proximal, distal, N3 and N4/N5 [27]. It is not currently known why MHV68 *Orf50* utilizes multiple different promoters or what the purpose of these various promoters are. It is possible that the different *Orf50* promoters have different functions, depending on cell type. One way to regulate the ability of these promoters to respond to various stimuli in different cells is to regulate the chromatin accessibility of the viral genome in macrophages and B cells. This differential chromosome accessibility may allow the virus to have distinct responses to various reactivation stimuli. The differential binding of STAT6 to the *Orf50* promoter suggests that the MHV68 genome displays different chromosome structure states depending on cell type.

MHV68 establishes latent infection in germinal center B cells and maintains long term latency in memory B cells [26,35]. T follicular cells make IL-4 as part of the germinal center reaction to promote class switching [33]. Therefore, even though germinal center and memory B cells that harbor latent virus encounter IL-4, they do not induce virus reactivation in response to IL-4 signals. This raises the question of whether the virus in latently infected B cells silences the *Orf50* N4/N5 promoter or preferentially uses different ORF50 promoters to regulate the latent to lytic switch.

The closely related human gammaherpesvirus, KSHV, also has 4 different *Orf50* promoters that induce expression of 6 different RTA transcripts [36]. There are also other herpesvirus open reading frames that are controlled by multiple promoters that produce alternative transcripts [37–39]. One possibility is that these different promoters respond to different acute, latent, and reactivation signals. Another possibility is that the different isoforms of RTA produced have different transactivation potential for various KSHV promoters. Both these possibilities reflect the intricate regulation of herpesvirus gene expression and illustrate the multiple ways the virus has evolved to regulate latency and reactivation.

During the lifetime of the host, the host and the latent virus will experience many bystander infections. We hypothesize that there are conditions which are favorable for virus reactivation and conditions which are unfavorable. Similarly, virus may reactivate from one tissue or cell type, but not from another tissue or cell type, depending on the signals received. The nature of chronic latent infection dictates that herpesviruses evolve exquisite sensitivity to the host environment.

## Author Contributions

GW, CZ, TAR designed experiments. GW and CZ performed experiments and analyzed data. GW, CZ, and TAR wrote and edited the manuscript. TAR secured funding. TC, LT, AL assisted with experiments.

## Acknowledgements

We thank the UTSW Flow Cytometry Core, the UTSW Animal Resource Center, UTSW IACUC for use of their facilities; the Ivan D’Orso lab for use of their sonicator; and Dr. Laurie Krug for providing HE2.1 B cells. This work was supported by the NIH (1R01AI130020-01A1), American Heart Association (17SDG33670071), CPRIT (RP200118), and the Pew Scholars Program.

---

**SI1.**
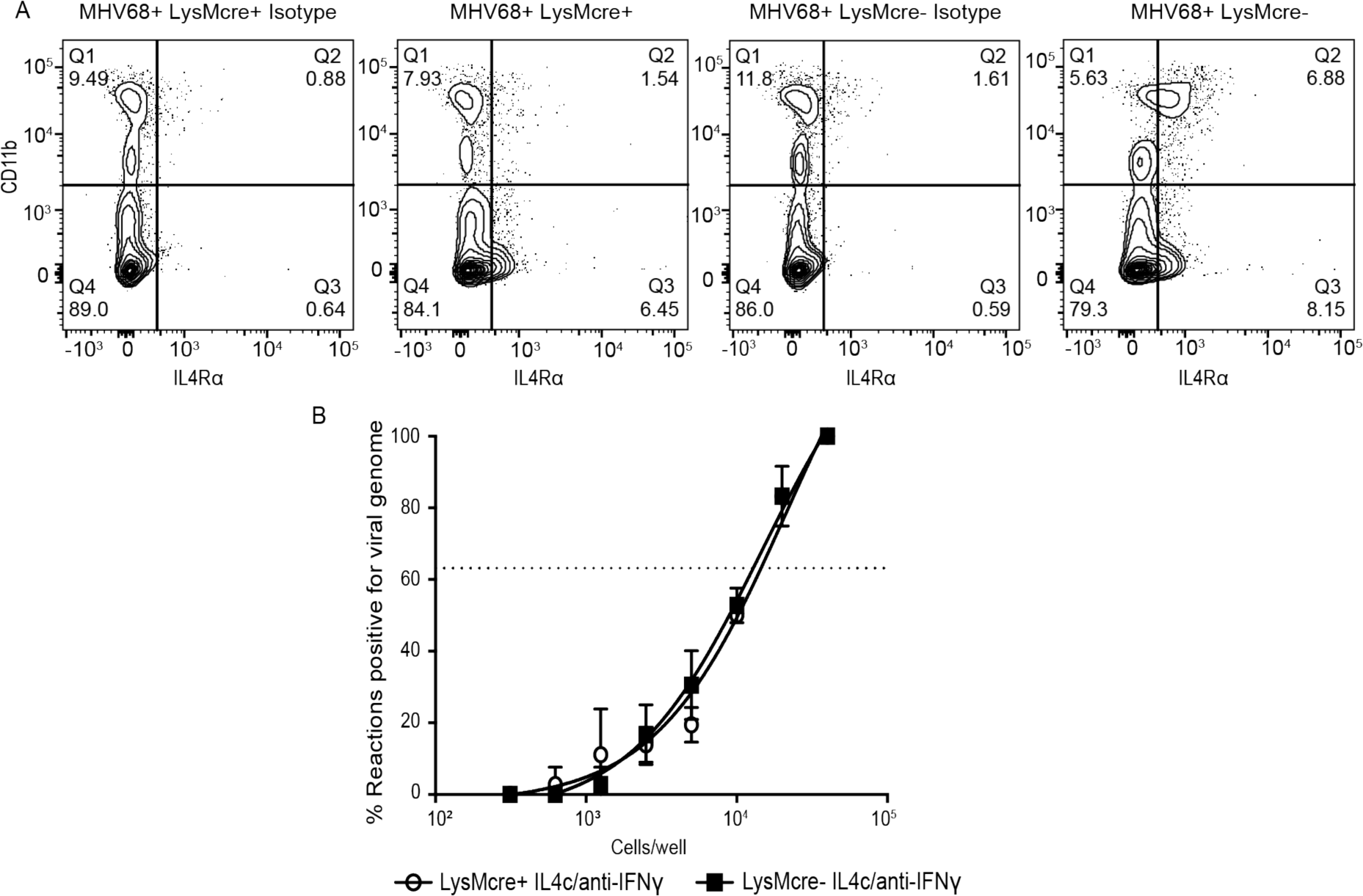
IL-4R expression is required for MHV68 reactivation from macrophages, Related to Figure 3. A. *Il4rα*^*f/f*^ *x LysMcre+* and *Il4rα*^*f/f*^ *x LysMcre-* mice were intraperitoneally infected with 10^6^ PFU wild-type MHV68. Peritoneal lavage was collected 37 days post infection, and flow cytometry was performed to detect IL-4R on peritoneal cells. Cells that have high expression of CD11b are macrophages and CD11b low or negative cells include B cells. Contour plots are representative of one experiment. B. Mice received both IL-4c and anti-IFNγ 28 days post-infection. Another dose of IL-4c was administered two days after the first treatment and then mice were euthanized at 5 days post-treatment. Peritoneal exudate cells were processed into single-cell suspensions and were subjected to nested-PCR analysis to assess the frequency of cells harboring viral genomes. Groups of 3-5 mice were pooled for each infection and analysis. The data are pooled from 3 independent experiments.

**SI2.**
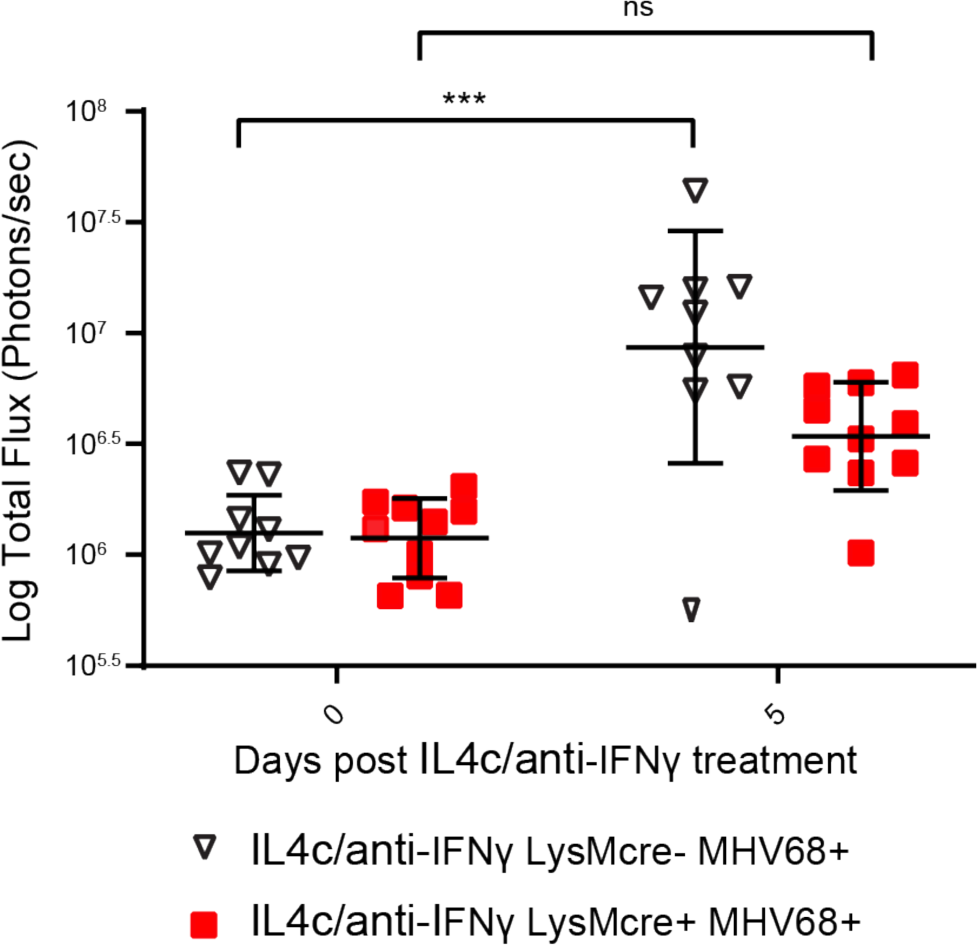
IL-4R expression is required for MHV68 reactivation from macrophages, Related to Figure 3. *Il4rα*^*f/f*^ *x LysMcre+* and *Il4rα*^*f/f*^ *x LysMcre-* mice were intraperitoneally infected with 10^6^ PFU MHV68-M3-FL. Mice received anti-IFNγ and IL-4c after 40 days of infection. IL-4c was administered two days after the first treatment, and then total flux (photons/second) from the abdominal region was quantitated at day 5 post-treatment using an IVIS bioluminescence imager. Bars represent mean ± SD. Each dot represents an individual mouse. The data are pooled from two independent experiments. ns, not significant, ***, P<0.001.

**SI3.**
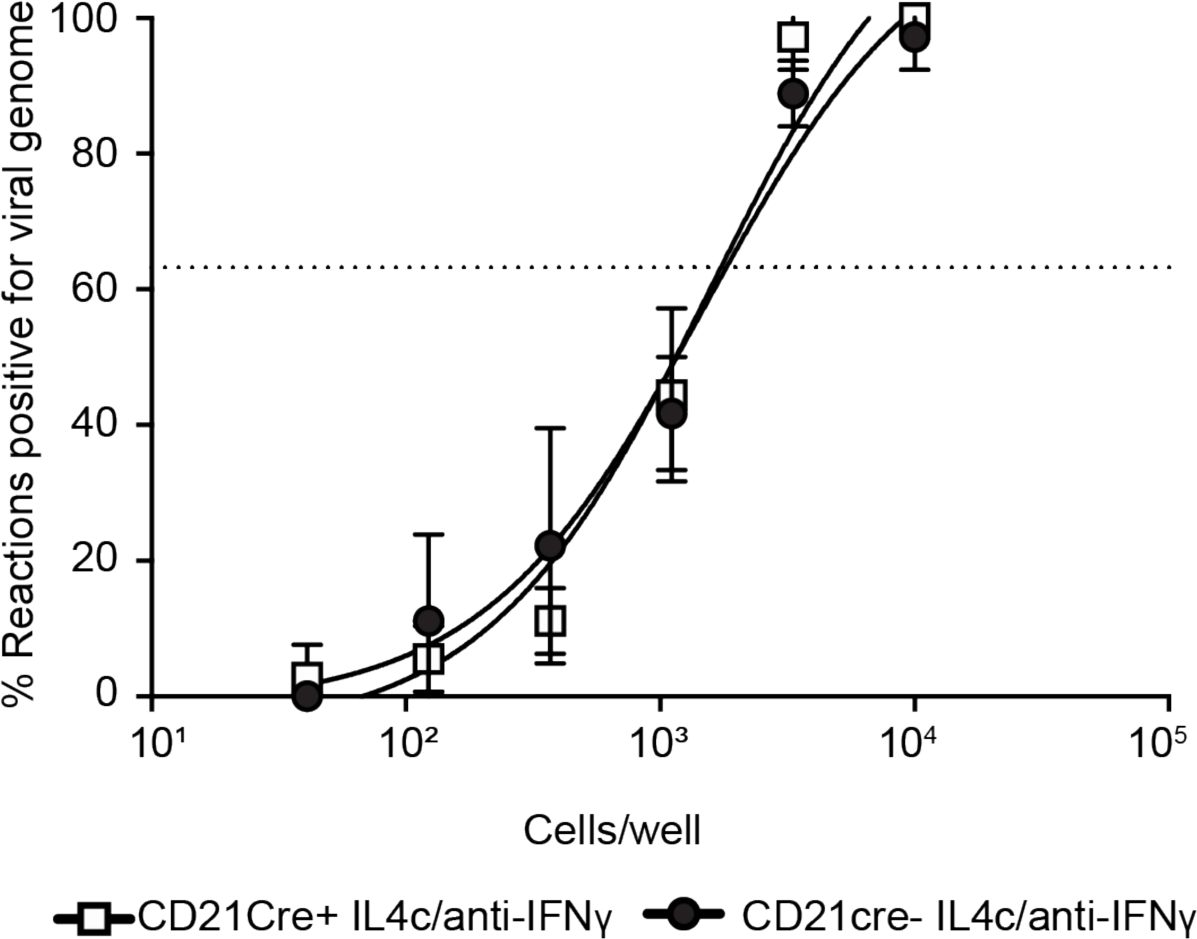
The absence of IL-4R on B cells does not alter establishment of MHV68 latency, Related to Figure 5. *Il4rα*^*f/f*^ *x Cd21-cre+* and *Il4rα*^*f/f*^ *x Cd21-cre-* mice were intraperitoneally infected with 10^6^ PFU wild-type MHV68. Mice received both anti-IFNγ and IL-4c after 28 days of infection. IL-4c was administered two days after the first treatment and then mice were euthanized at 5 days post-treatment. Peritoneal exudate cells were processed into single-cell suspensions and were subjected to nested-PCR analysis to assess the frequency of cells harboring viral genomes. Groups of 3–5 mice were pooled for each infection and analysis. The data are pooled from 3 independent experiments.

## MATERIALS AND METHODS

### Animals, infections and treatment

All mice used in this study were from C57BL/6 background and housed in a specific pathogen-free facility at the University of Texas Southwestern Medical Center. Mice were maintained and used under protocol approved by UT Southwestern Medical Center Institutional Animal Care and Use Committee (IACUC). To generate mice conditionally deficient for *IL4r, IL4raf/f* mice [16] were crossed to Lysozyme M-cre recombinase expressing mice [40] or CD21-cre recombinase [41] expressing mice for selective disruption of *IL4ra* in different cell types *in vivo*. Mice were used for infections between 6-12 weeks of age and all experiments used littermate controls. Mixed groups of both males and females are used in all experiments. MHV68 (WUSM strain) and MHV68-M3-FL [20] diluted in D10 (DMEM with 10% FBS) and were administered intraperitoneally (10^6^ PFU).

### *In vivo* MHV68 reactivation

For IL-4c/anti-IFNγ induced reactivation, both anti-IFNγ and long-lasting IL-4 complexes (IL-4c) were injected intraperitoneally on day 28 of MHV68 infection and a second dose of IL-4c was injected on day 30 as described in Reese et al [7]. 500 μg of anti-IFNγ (BioXCell, R4-6A2) or isotype control (BioXcell, clone HRPN) was diluted in PBS. 5 μg of IL-4 (Peprotech, AF-214-14) and 25 μg of anti-IL-4 (BioXcell, clone 11B11) were complexed prior to intraperitoneally injection into mice. For cyclosporine A-induced reactivation, 50 mg/kg of cyclosporine A (Sigma, lots 098M4084V and 095M4084V) with corn oil as a carrier were injected intraperitoneally into the infected mice on the days indicated on figures. Mice were imaged on day 0 before cyclosporine A injection, and days 5, 7, 9, and 12 following treatment.

### Cell cultures and infections

Murine fibroblast (3T12, ATCC CCL-164) cells were cultured in Dulbecco’s modified Eagle medium (DMEM; Corning) supplemented with 5% fetal bovine serum (Biowest), 2 mM L-glutamine (Gibco), 1% HEPES (Corning), 10 U/ml penicillin, and 10 μg/ml streptomycin sulfate (Corning) (complete DMEM). Cells were cultured at 37°C with 5% CO2. A20-HE2.1 B cell lines [14] (kindly provided by Laurie Krug) were maintained in RPMI (Gibco), 10% fetal bovine serum, 2 mM L-glutamine, 100 U/ml penicillin, 100 mg/ml streptomycin, 50 μM *β*-mercaptoethanol and 300 µg/ml G418. Mouse embryonic fibroblasts (MEFs) were obtained from C57BL/6 mouse embryo. BMDMs were harvested from C57BL6/J mice and differentiated for 7 days with 10% CMG-14 supernatants [42] in complete Dulbecco’s modified Eagle’s medium (DMEM) (10% fetal bovine serum, 1% HEPES, 2mM L-glutamine). At day 7, 5×10^5^ cells per well were plated in 6-well plates, pretreated with IL-4 (Peprotech) for 16 hours, infected with MHV68 one hour and then washed with PBS and cultured in media containing cytokines.

### Generation of MHV68 stock

Viral stocks were prepared by low MOI infection of 3T12 cells in 5% FBS complete medium. Virus stocks were harvested after 7 days of infection and were processed by freeze/thawed cycles. Subsequently, cell debris was pelleted by centrifugation at 3000 rpm for 30 minutes, and purified virus stock was prepared by centrifugation at 8000g for 2 hours at 4°C, and aliquots were transferred to −80°C for long-term storage. Concentration of virus stocks were determined by plaque assay using on 3T12 cells.

### Determination of MHV68 viral titer

3T12 cells were seeded in 6-well plates at 3×10^5^ cells/well. Homogenized tissue or viral stock supernatants were serially diluted in D5 medium (DMEM, 5% FBS, 2 mM L-glutamine, 1 mM HEPES) and added to 3T12 cells monolayers for 1 hour at 37°C. Samples were overlaid with 1% methylcellulose in DMEM with 5% FBS. Plates were incubated at 37°C for 7 days, and then stained with crystal violet (0.2% crystal violet in 20% ethanol) to visualize plaques.

### Ex vivo limiting dilution assay (LDA) for reactivation and persistent replication

To determine the frequency of cells harboring latent virus capable of reactivation *ex vivo*, single cell suspensions from peritoneal exudate cells (PECs) and spleens were plated in two-fold serial dilutions on a mouse embryonic fibroblast (MEF) monolayers (maintained in DMEM with 10% FBS) [18]. Cytopathic effect (CPE) was scored three weeks after plating. To distinguish preformed infectious virus from virus that reactivates ex vivo from live cells upon explantation, parallel samples were mechanically disrupted using 0.5-mm silica beads on the Precellys 24 tissue homogenizer (Bertin Technologies) to kill the cells, but keep any infectious virus intact and then plated on the monolayer of MEFs to release preformed virus. The 63.2% Poisson distribution horizontal line represents the frequency at which one reactivation event is likely to occur per number of plated cells.

### Limiting Dilution (LD)-PCR to detect viral genome

To compare the frequency of cells the harboring viral genome, single cell suspensions from PECs and spleens were assayed by nested PCR [5]. In brief, cells were serially diluted in a solution containing uninfected 3T12 cells to maintain a total number of 10^4^ cells per PCR reaction and lysed overnight at 56°C with proteinase K. Two rounds of PCR were performed using six dilutions per sample with 12 reactions per dilution with primers specific for MHV68 gene ORF72. Plasmid containing ORF72 was included at 0.1, 1.0, and 10 copies as positive controls. Products were visualized on 1.5% agarose gel. Data points for LD-PCR represent the mean and the standard error of the mean for three replicate experiments. Frequencies of viral genome-carrying cells were obtained by calculating the cell density at which 63.2% of the wells were positive based on the Poisson distribution.

### Bioluminescence imaging of mice using MHV68-M3-FL

Mice were infected with 10^6^ PFU per mouse with MHV68-M3-FL. The mice were anesthetized in isoflurane and imaged after intraperitoneal injection of 150 mg/kg of D-luciferin (Gold Biotechnology). Bioluminescence imaging of MHV68-M3-FL infected mice was collected on the IVIS Lumina Series III (Perkin Elmer), and data collection and analysis were done on the Living Image software (Caliper Life Sciences). Mice were exposed for 3 min, for 3 consecutive pictures. Total luminescence was calculated by measuring total flux (photons/sec) in a region of interest (ROI) for each mouse. One ROI size was used for each image, to accommodate the size of the mice. The pictures with peak luminescence were selected as final data.

### Flow cytometry

Whole spleens were processed into a single cell suspension and strained. Peritoneal exudate cells were collected by peritoneal lavage and washed with PBS. Any samples with blood were lysed with ACK lysis buffer (Lonza BioWhittaker). Fc receptors were blocked on cells with α-CD16/32 (2.4G2, Tonbo). Cells were stained with the following: BV605-α-CD19 (1D3, BD Biosciences), APC-Cy7-α-CD19 (1D3, BD Biosciences), Per-CpCy5.5-α-F4/80 (BM8.1 Tonbo), APC-Cy7-α-F4/80 (BM8, Biolegend), Fitc-α-CD11b (M1/70, Biolegend), PE-Cy7-α-CD11b (M1/70, Tonbo), PE-α-IL4Rα/CD124 (I015F8, Biolegend), Violetfluor 450-α-CD3 (17A2, Tonbo). Rat PE-α-IgG2b (LTF-2, Tonbo) was used as the isotype control for CD124. Cells were suspended in FACS buffer (PBS with 1% FBS) and then analyzed on an LSR II flow cytometer. Data was analyzed using FlowJo software (TreeStar, Inc., San Carols, CA).

### Real-time qPCR

RNA was isolated using RNeasy mini kit (Qiagen) and cDNA was prepared using SuperScript VILO cDNA Synthesis Kit (Invitrogen) according to manufacturer’s instruction. Quantitative PCR was performed using PowerUp SYBR Green Master Mix (Applied Biosystems) in a Quant Studio 7 Flex real time PCR system using the primers listed in Table 1. QPCR is performed on three biological replicates and fold change was calculated by ΔΔCT method.

**Table 1.**
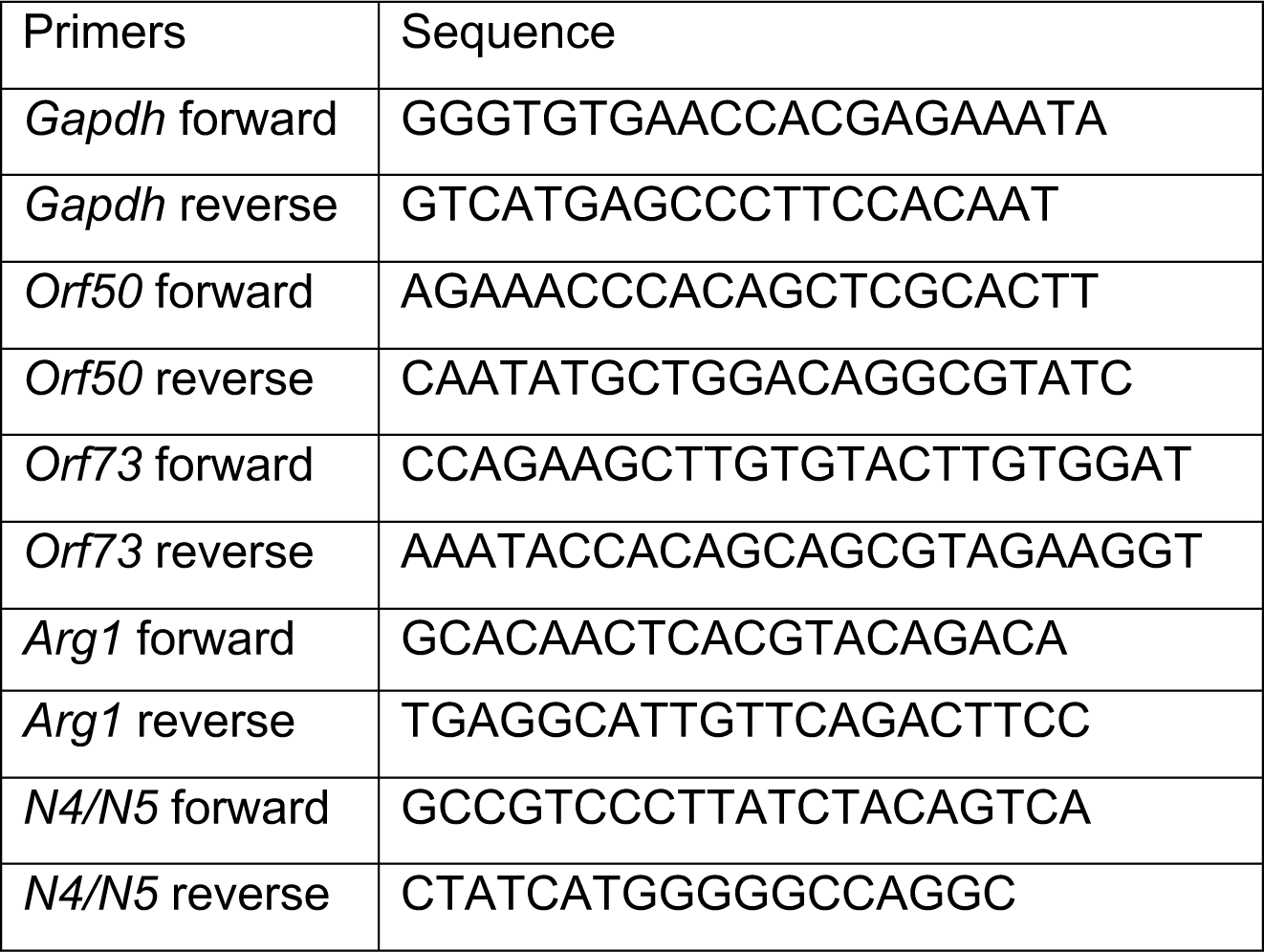

### Immunoblot analysis

Total protein lysate was harvested in RIPA buffer (150 mM NaCl, 1% NP-40, 0.5% sodium deoxycholate, 0.1% SDS, 25 mM Tris with protease inhibitor cocktail). Protein concentration were determined using Bradford assay (Bio-Rad). Proteins were separated on 4-12% Bis-Tris plus gels (Thermo Fisher Scientific) and transferred to nitrocellulose membrane. Antibodies against STAT6 (Santa Cruz Biotechnology, Dallas, TX), tyrosine 641-phosphorylated STAT6 (Santa Cruz Biotechnology, Dallas, TX), and actin (1:5000, Catalog no. A2228, Sigma) were detected by the use of secondary donkey-anti-rabbit (1:5000, Catalog no.711-035-152, Jackson Immuno Research Laboratory) and goat-anti-mouse peroxidase (1:5000, Catalog no.115-035-174 Jackson Immuno Research Laboratory) by immunoblot analysis with Luminata Forte Western HRP substrate (Millipore).

### Chromatin immunoprecipitation

Cells from a 10 cm dish were crosslinked in 1% formaldehyde in PBS for 10 minutes at room temperature, quenched in 0.125 M glycine for 10 minutes. The crosslinked cell suspension was then spun down at 800 *g* at 4 °C for 5 minutes. The cell pellet was washed twice with cold PBS and lysed with 1 ml ChIP lysis buffer (50 mM HEPES pH 7.9, 140 mM NaCl, 1 mM EDTA, 10% glycerol, 0.5% NP40, 0.25% Triton X-100). Nuclei were collected by centrifugation at 1000 g for 10 minutes at 4°C and resuspended in 0.5 ml of ChIP shearing buffer. Samples were then sonicated until DNA was fragmented to an average distribution of about 200–300 bp using a Bioruptor (Diagenode) for a total of 30 cycles (30s on and 30s off). Sonicated nuclear lysate was spun at 15,000 g for 5 minutes at 4°C and the pellet was discarded. Sonicated nuclear lysate was incubated with 10 µg rabbit monoclonal anti-Stat6 (sc-374021) or rabbit IgG overnight. 20 µl of mixed protein A and G dynabeads (Thermo Fisher) were added and tubes were rotated for 2 hours at 4 °C. Beads were washed with low salt immune complex (20 mM Tris pH 8.0, 1% NP-40, 2 mM EDTA, 150 mM NaCl, 0.1% SDS, 0.5% deoxycholic acid), high salt immune complex (20 mM Tris pH 8.0, 1% NP-40, 2 mM EDTA, 500 mM NaCl, 0.1% SDS, 0.5% deoxycholic acid), lithium chloride immune complex (10 mM Tris pH 8.0, 0.25 M LiCl, 1% NP-40, 1% deoxycholic acid, 1 mM EDTA), and Tris-EDTA twice. After washing, complexes were eluted off the beads using elution buffer (1% SDS and 100 mM NaHCO3) at 65 °C for 30 minutes. Protein-DNA complexes were de-crosslinked using de-crosslinking buffer (50 mM Tris–HCl pH 6.8, 500 mM NaCl, 5 mM EDTA, 0.5 mg/mL Proteinase K) at 60 °C overnight. DNA was then purified using the Zymo ChIP DNA clean and concentrator kit (Zymo Research) and used for qPCR analysis using the primers listed in Table 1.

### Statistical analysis

Data were analyzed with Prism 7 software (GraphPad Software, San Diego, CA). In all graphs, significant value was indicated as; ∗, p < 0.05; ∗∗, p < 0.01; ∗∗∗, p < 0.001; which determined by one-way analysis of variation (One-way ANOVA) with multiple comparisons (for multiple samples), Student’s t test (for two samples) as indicated in figure legends.

